# Assessing nasal obstruction with the nasal acoustic device: a pilot study on distinguishing between subjects with and without chronic rhinosinusitis

**DOI:** 10.1101/631499

**Authors:** Chia-Hung Li, Calvin Tan, Katherine L. Whitcroft, Peter Andrews, Terence S. Leung

## Abstract

This article aims to demonstrate the concept and potential of a novel diagnostic device – the nasal acoustic device (NAD), which captures the nasal breathing sounds over the ala on both sides of the nose. Using the newly defined inspiratory nasal acoustic score (INA score), the unique characteristics of the nasal breathing sounds can be quantified and used for diagnostic purposes.

In this pilot study, the NAD was compared with the well-established nasal inspiratory peak flow meter (NIPF) to try to distinguish between subjects with and without chronic rhinosinusitis (CRS). Measurements were made before and after nasal decongestants, which were applied to eliminate the nasal cycle. Patients were divided into two groups based on the presence or absence of CRS, irrespective of other pathological conditions being present.

Based on the post-decongestant measurements, the sensitivity/specificity in distinguishing subjects who have CRS (n=11) from subjects who do not (n=29; of whom 13 were controls) were 70%/66% for the NIPF and 82%/79% for the NAD. The non-CRS groups showed statistically significant changes after decongestant for both methods, but the CRS group did not. The results firstly demonstrated that the CRS subjects in this cohort tended to be less responsive to decongestant and therefore the post-decongestant NIPF measurements provided a certain degree of diagnostic value in identifying CRS subjects, but it would appear the NAD better captured the unique sounds associated with CRS, providing a superior diagnostic capability. This study also demonstrates that the NAD can measure the improvement in the nasal airway following treatment effect.

## Introduction

Nasal obstruction has a prevalence rate of over 30% amongst the general adult population [1]. Generally defined as a feeling of discomfort due to inadequate nasal airflow, it is commonly caused by two inflammatory disorders: allergic rhinitis (AR) and chronic rhinosinusitis (CRS). The prevalence rate of AR in Europe is around 25% [2] and for CRS it is around 11% [3]. Nasal obstruction can also be caused, albeit less commonly, by a structural blockage as opposed to an inflammatory swelling, such as a deviated nasal septum (DNS). As a result nasal obstruction is one of commonest complaints relating to ear, nose and throat (ENT) in primary care and one of the commonest reasons for secondary care referrals for ENT [4, 5].

According to our survey of ENT practitioners, the majority of ENT surgeons diagnose nasal obstructions using only the patient’s history and nasal endoscopy; with only a few using a medical device such as the nasal inspiratory peak flow (NIPF) meter [6]. GPs in primary care also predominantly use history and examination, the latter with a simple pen torch rather than nasal endoscopy. Current evidence shows that patients with persistent nasal obstruction in the UK can wait up to 5 years before their conditions are diagnosed and treated by secondary care clinicians [7].

There is increasing evidence which demonstrates that delayed surgery for persistent CRS, which is not responding to maximal medication, can lead to long term negative effects for the patient as a result of potential irreversible damage to the nasal lining. Patient’s with delayed surgical intervention demonstrate a reduced post-operative quality life (QOL) [8] and increased post-operative health care needs [9]. In this study CRS patients would require significantly more CRS-related care if they underwent surgery after 5 years of diagnosis as opposed to having it within 1 year; demonstrating that irreversible damage could occur if the treatment is delayed [9]. In addition CRS patients who end up with a delay in their medical treatment demonstrate a reduced olfactory improvement compared to those whom were treated earlier in their CRS journey [10].

The unmet need is a medical device which could help in diagnosing the cause of nasal blockage and help reduce the delay in treatment as well as reduce the need for secondary care referral [11]. The three commonly used medical devices which are used to assess the function of the nose are; the nasal inspiratory peak flow meter (NIPF) which measures the nasal airflow, acoustic rhinometry, which measures the minimal cross-sectional area of the nose, and rhinomanometry which measures the nasal airway resistance. NIPF is the most commonly used device according to our survey because it is inexpensive, portable as well as quick and easy to use [12]. It has undergone multiple validation studies to determine its normal range [13–16]. Until recently the NIPF has only been used to measure bilateral nasal airflow which has made difficult to assess unilateral obstruction. However, the device is now being validated for unilateral measurement, by covering one nostril before measurements [17].

Other medical devices also exist but are mainly used for research purposes. One example is nasal spirometry which measures the volume of air expired from each nostril [18]. Another method is the use of polyvinylidene fluoride (PVDF) strip directly underneath each nostril to measure the nasal airflow [19]. Pressure sensors are also used in the case of the nasal data logger to monitor the nasal cycle [20]. The Glatzel mirror allows straight forward observation of the condensation pattern created when air is expired on it [21]. A video device, the video rhino-hygrometer, was developed to attempt to quantify such a misting pattern [22].

Acoustic sensors have been previously explored by placing one or more microphones underneath the nostrils or somewhere along the direct path of the nasal airflow and the signals are generated when expired nasal air hits the microphones [23]. These studies include the Odiosoft-Rhino (OR) technique [24], the nasal sound spectral analysis (NSSA) [25] and the Primov-Fever *et al* description [26]. These studies share one common overarching concept which is to measure the nasal airflow through direct interaction of the sensor with airflow. While most non-acoustic methods also follow this principle, in the case of acoustic this unfortunately can cause distortions.

The general aim of this pilot study is to evaluate the nasal acoustic device (NAD) which is based on a novel ‘stethoscope’ concept whereby a microphone is placed over each nasal ala so that airflow cannot directly interact with the sensors and in doing so avoiding the aforementioned distortions. The nasal breathing sounds detected are caused by air turbulence generated by the internal structure and influenced by tissue characteristics of the nose during both nasal inspiration and expiration [26].

In this pilot study we aim to assess the diagnostic accuracy of the NAD in detecting CRS from the other common causes of nasal blockage which include AR and DNS by comparing its performance with NIPF. This study had demonstrated the potential diagnostic capability in detecting the presence of CRS from a relatively diverse subject groups composing of both controls and patients with a combination of two different conditions which include AR and DNS.

## Materials and Methods

A study was carried out at a specialist ENT hospital, with ethical approval by London - City & East Research Ethics Committee. Written consent forms were provided for the participants. Both patients with nasal obstructions and control subjects were recruited. Patients were recruited if they had at least one of: allergic rhinitis, chronic rhinosinusitis, or deviated nasal septum. The exclusion criteria were if the subjects had systemic diseases involving the nose such as sarcoid or vasculitis; had nasal tumours; or were below 18 years old. A patient’s condition was diagnosed using nasal endoscopic examination by a consultant ENT surgeon (co-author PA). Control subjects were University College London students who were recruited and assessed in the same manner as the patients. All subjects took part in the study at the Royal National Throat, Nose and Ear Hospital. Each subject was assigned to one of two groups: the CRS group or the Non-CRS group, based on whether he or she has CRS and irrespective of whether the subject had any other conditions. This means that subjects in the two groups can be similar in that they can all have allergic rhinitis and/or DNS, but only the CRS group will have CRS. The non-CRS group was also sub-divided into controls and patients without CRS.

### Method of Study

NIPF and nasal breathing sound were measured for each patient prior to and 10 minutes after applying nasal decongestant. For each method the post-decongestant percentage change in the measured parameters were calculated.

The decongestant used was a mixture containing: the actual decongestant agent (phenylephrine hydrochloride 0.5% w/v); local anaesthetic to numb the airways (lidocaine hydrochloride 5% w/v); plus a preservative (benzalkonium chloride). These were mixed in distilled water. It was sprayed twice or thrice into each nostril.

### Nasal inspiratory Peak Flow Meter

NIPF measurements were performed with the patient in a sitting position. The patient was asked to take a deep breath before covering the nose and mouth area with the mask of the meter. The patient then made a single inspiration through the nose, at the maximum effort and speed, with his or her mouth closed. NIPF was performed 3 times per patients, the largest of which was used for analysis. Measurements were repeated if one of the three readings deviated from the others by 40 L/min or above.

### Nasal Acoustic Device

Figure 1 shows a diagram for the nasal acoustic device and Figure 2 shows an example acoustic recording. A modified, contact-based piezo-electric microphone (Nordell Acoustic Guitar Pickup, Dangleberry Music, UK) was placed each side of the nose (i.e. one per side). A strip of bio-compatible double-sided tape (Body Tape, Eylure, UK) was used to adhere the microphone onto the nose. Additional porous tapes (Micropore Surgical Tape, 3M, USA) were used to adhere the wires of the microphones onto the face to minimise the chance of them falling off.

**Fig.1.**
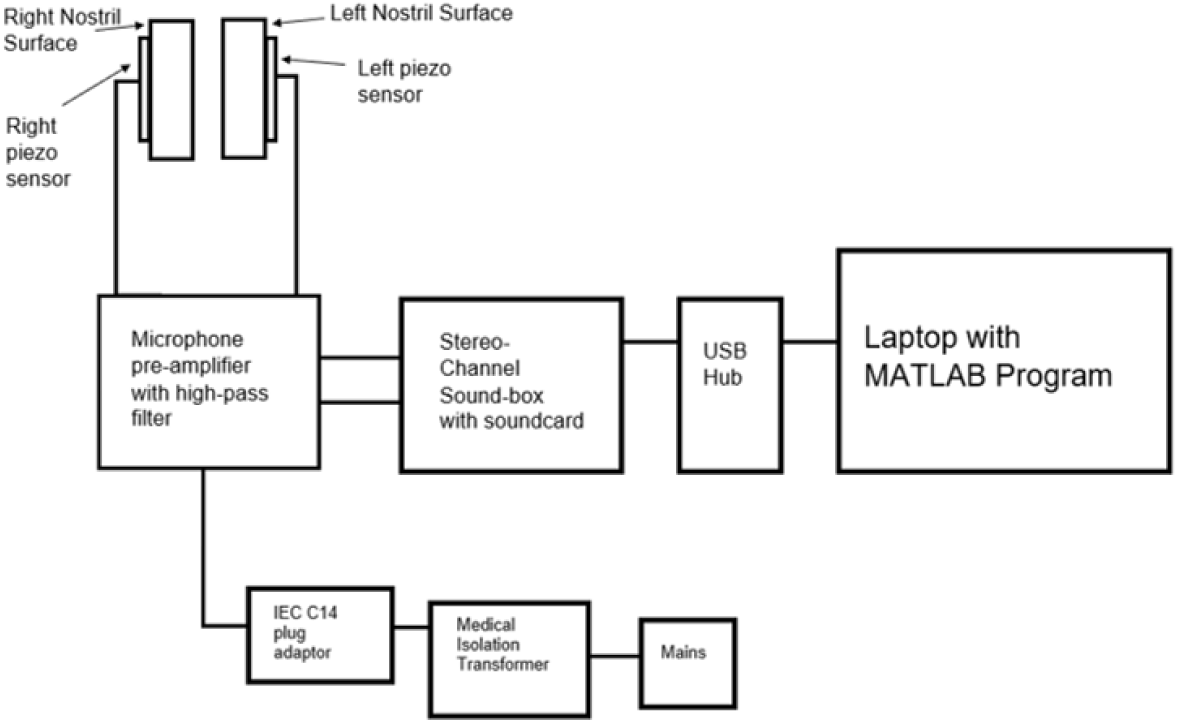
Block diagram of components for NAD.

**Fig.2.**
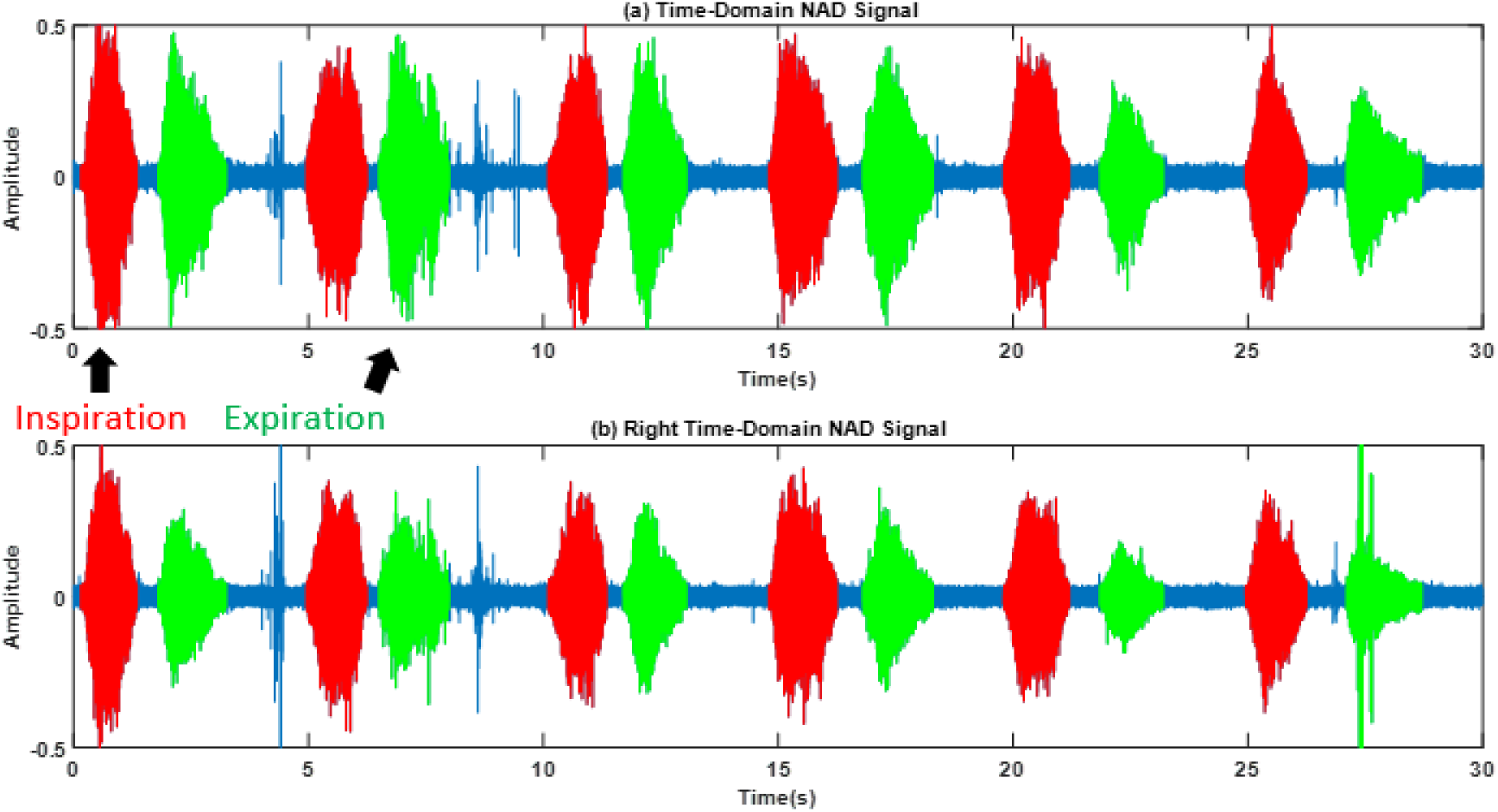
An example of time-domain acoustic signals recorded using NAD, with left (a) and right (b) sides recorded unilaterally and simultaneously. The inspirations and expirations as detected by the program were shown in red and green respectively. Any parts of the signal that remains blue were considered background noise.

The microphones were connected to a two-channel amplifier (Stage Line MPA-202, Monacor International, Germany) with built in higher-pass filters. The amplification was set to 65dBa amplification and the cut-off frequency was set to 60Hz. Each channel of the amplifier output was connected to a stereo sound card (SW-29545, Sewell Direct, USA) which in turn was connected via an USB channel to a laptop (Dell Inspiron 5567, Dell, USA). Acoustic signals were acquired using a MATLAB (MATLAB R2018a, MathWorks, USA) program. The sampling frequency was set to 44,100Hz. Both the laptop and the amplifier were connected to the mains power supply via a medical grade isolation transformer (REOMED 200-230V, REO AG, Germany). The system was tested and certified for electrical safety by the University College London Hospital.

Each acoustic measurement was performed with the subject in a sitting position. Breathing sound was recorded over a 30 second period during which the patient was asked to make light-to-moderate effort breathing throughout, with the mouth closed. Breathing rhythm was kept as natural as possible (e.g. a patient should not make a very deep and lengthy nasal inspiration before proceeding to give a small and short expiration on purpose). The NAD operator can demonstrate for the patients the desired breathing effort

### Data Analysis

A separate MATLAB program was used to analyse the acoustic signals. Prior to analysis, the acoustic signals were furthered bandpass-filtered at 100Hz-20kHz by the program. The program can detect nasal inspirations and expirations as well as using them to calculate various metrics that may be of diagnostic value. The method for detecting inspirations and expirations is beyond the scope of this article, however the detected breathing phases were verified by listening to .wav files created for the recordings. In addition, the program rejects sudden increase in background noise that may appear to be breathing sounds (as seen in the example of Fig.2.).

This study presents a novel inspiratory nasal acoustic (INA) score which is shown in Equation 1. The INA score is the average acoustic power of the nasal inspiration at a specific frequency, expressed as a percentage of the acoustic power for the entire frequency spectrum. A frequency band of 600Hz to 7900Hz was chosen using a third MATLAB test program that was used to systematically identify the optimal frequency band. The optimal frequency band was chosen based on the Youden number (see the statistical Analysis section): a number that shows the optimal sensitivity and specificity in distinguishing between the two groups of subjects. One rationale for normalizing the acoustic power was to account for the fact that different patients may breathe at different efforts. For each recording, the average band-power of all inspiratory phases was calculated. Because CRS is a bilateral condition and the NIPF measured for this study was also bilateral, the NAD would also be using bilateral results. Since NAD records breathing sound from each nostril unilaterally, the measurements for the two sides were averaged out to give an overall bilateral reading, which was used to calculate the INA score.

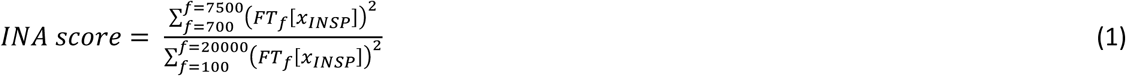

Where f = frequency and x_INSP_ represents the time-domain signals in the inspiratory phase.

### Statistical Analysis

The post-decongestant percentage changes of the measured values were calculated. For each subject group and each measurement method, the percentage change was compared with zero using Wilcoxon signed-rank test.

Receiver operating characteristic (ROC) curves were generated for both methods for post-decongestant measurements. The optimal cut-off values to differentiate between CRS and non-CRS groups were found using the principle of Youden’s number [27] and the corresponding sensitivity and specificity values as well as the positive and negative predictive values (PPV and NPV) were calculated. The area under curve (AUC) for each ROC curve was also found.

## Results

### Patient Characteristics

Table 1 displays the demographics of the subjects based on their conditions. A total of 27 patients (mean age 33±9; 15 males and 12 females) and 13 controls (23±2, with 10 males and 3 females) took part in the study. The non-CRS patient group had n=16 patients, which along with the n=13 controls, gives a total of n=29 non-CRS subjects; while the CRS patient group had n=11 patients. One patient did not perform pre-decongestant NIPF measurements and another patient did not perform post-decongestant NIPF measurements.

**Table 1:** The number of subjects for each condition (or combination of conditions).

| **Non-CRS subjects** | **n** | **CRS patients** | **n** |
| --- | --- | --- | --- |
| Controls | 13 | CRS only | 2 |
| Allergic rhinitis only | 2 | CRS + allergic rhinitis | 1 |
| DNS only | 9 | CRS + DNS | 2 |
| Allergic rhinitis + DNS | 5 | CRS + allergic rhinitis + DNS | 6 |
| Total | 29 | Total | 11 |

### Data Analysis

Figure 3 shows the boxplot for the pre- and post-decongestant measurements in NIPF for CRS and non-CRS subjects; while Figure 4 shows likewise for INA score.

**Fig.3.**
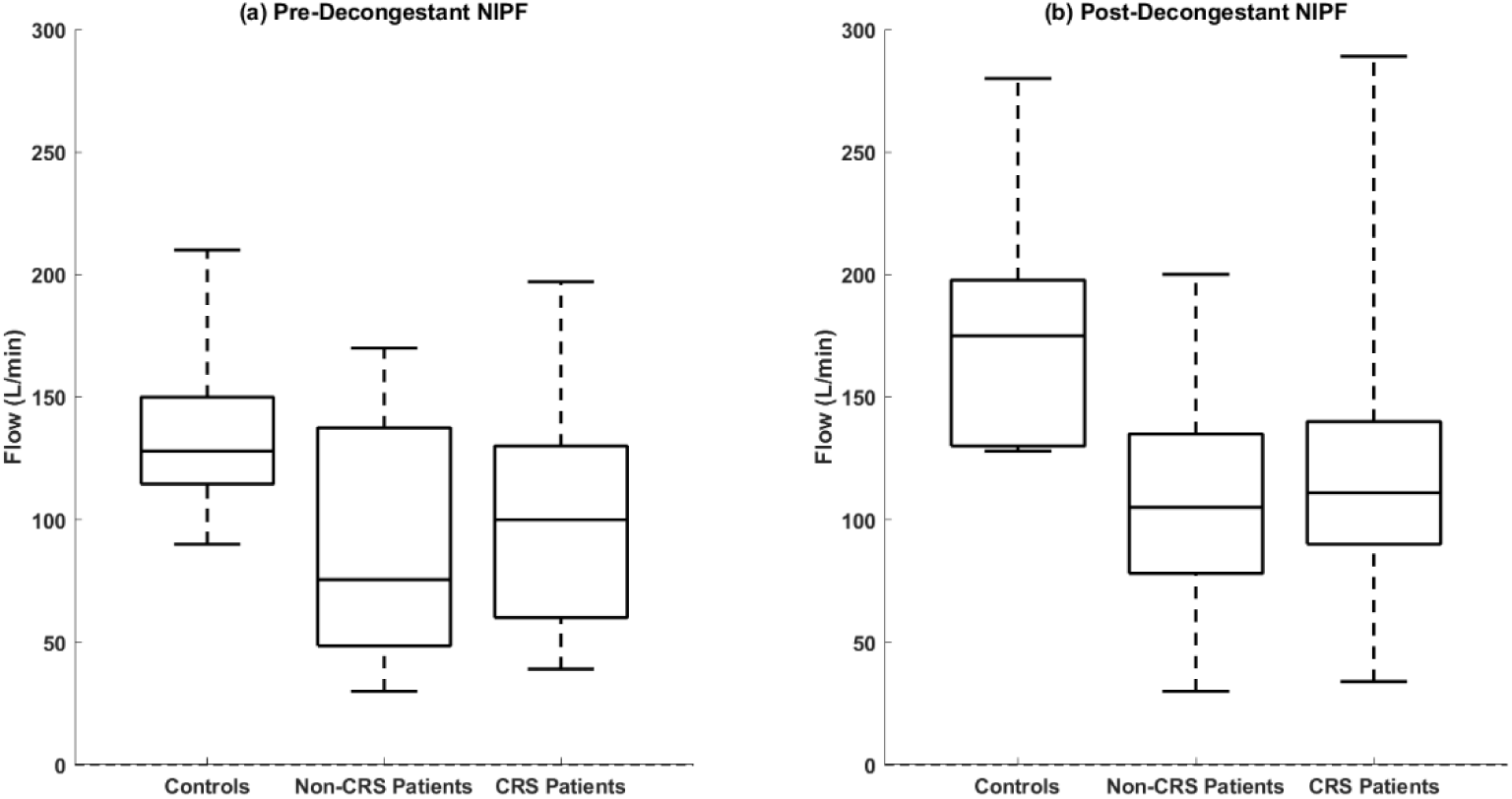
Boxplots display for the (a) pre- and (b) post-decongestant measurements in NIPF.

**Fig.4.**
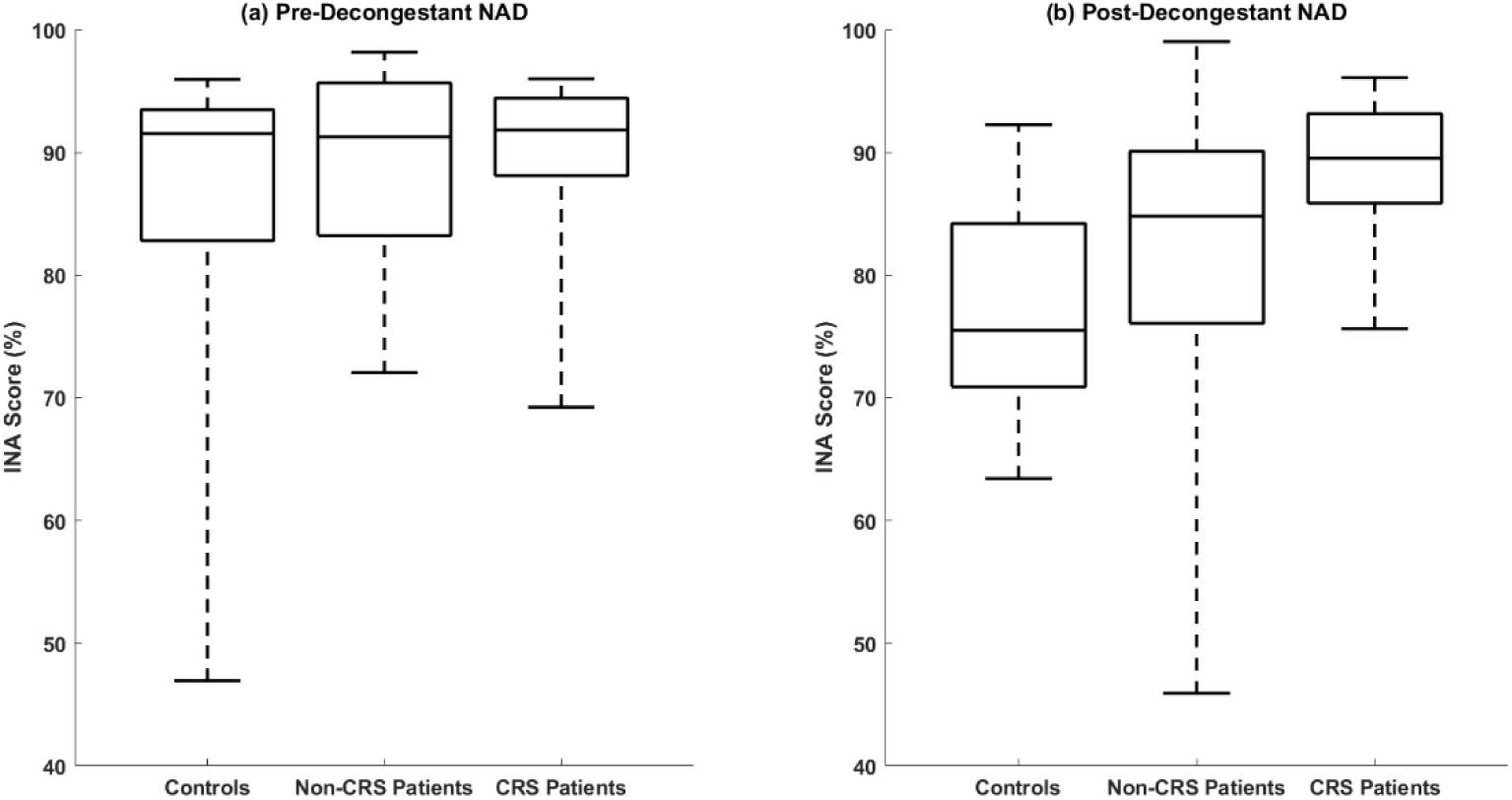
Boxplots display for the (a) pre- and (b) post-decongestant measurements in NAD.

Table 2 displays the median and range for all measurements and all groups as well as the post-decongestant percentage changes. The post-decongestant percentage changes were statistically significant for controls and non-CRS patients, be it NIPF or INA score, as shown when comparing their values with zero (p<0.05 in all cases). On the other hand, the change was not significant for the CRS patients (p>0.05) for either methods. The existence of outlier values in control and non-CRS patient groups meant that their percentage change in INA score appeared closer than they otherwise would have been.

**Table 2:** Median and range (in brackets) of NIPF and INA score for controls, non-CRS patients, and CRS patients; pre- and post-decongestant along with post-decongestant percentage changes. Percentage changes were compared with zero using Wilcoxon signed-rank test, with statistically significant results (p<0.05) in bold.

|  | NIPF (L/min) |  |  | INA score (%) |  |  |
| --- | --- | --- | --- | --- | --- | --- |
|  | Controls | Non-CRS Patients | CRS Patients | Controls | Non-CRS Patients | CRS Patients |
| Pre-Decongestant | 128.0<br>(90.0 to 210.0) | 75.5<br>(30.0 to 170.0) | 100.0<br>(39.0 to 197.0) | 88.3<br>(43.5 to 94.4) | 89.4<br>(67.4 to 97.4) | 90.1<br>(62.5 to 94.7) |
| Post-Decongestant | 175.0<br>(128.0 to 280.0) | 105.0<br>(30.0 to 200.0) | 111.0<br>(34.0 to 289.0) | 73.3<br>(61.3 to 90.7) | 81.8<br>(48.7 to 98.3) | 88.8<br>(69.5 to 95.3) |
| % Change | 33.3 %<br>(-27.8 to 81.8) | 30.6 %<br>(-50.0 to 433.3) | 12.5 %<br>(-28 to 150) | -12.6 %<br>(-24.0 to 58.2) | -9.8 %<br>(-33.7 to 17.5) | -1.2 %<br>(-14.4 to 52.5) |
| Statistically significant % change? | **p = 0.004** | **p = 0.019** | p = 0.102 | **p = 0.020** | **p = 0.046** | p = 0.289 |
|  | Increase after decongestant |  |  | Decrease after decongestant |  |  |

Figure 5 displays the ROC curves for NIPF (Fig.4a.) and INA score (Fig.4b.) respectively in distinguishing CRS patients from the non-CRS subjects (controls and patients together) using only the post-decongestant measurements. Table 3. displays the results of statistical comparisons between CRS patients and non-CRS subjects. When classifying between CRS and non-CRS subjects, using a cut-off of 126 L/min, the sensitivity/specificity for NIPF was 70%/38% when control subjects were not included, and 70%/66% if they were. Likewise, the sensitivity/specificity for INA score was 82%/69% without the control subjects, and 82%/79% with them; using a cut-off threshold of 84%. PPV/NPV for NIPF were 41%/83% regardless of whether controls were included. For INA score PPV/NPV were 64%/83% without controls and 60%/91% with controls. AUC for NIPF was 0.446 without controls and 0.617 with controls; and for INA score it was 0.665 without controls and 0.757 with controls.

**Table 3:**
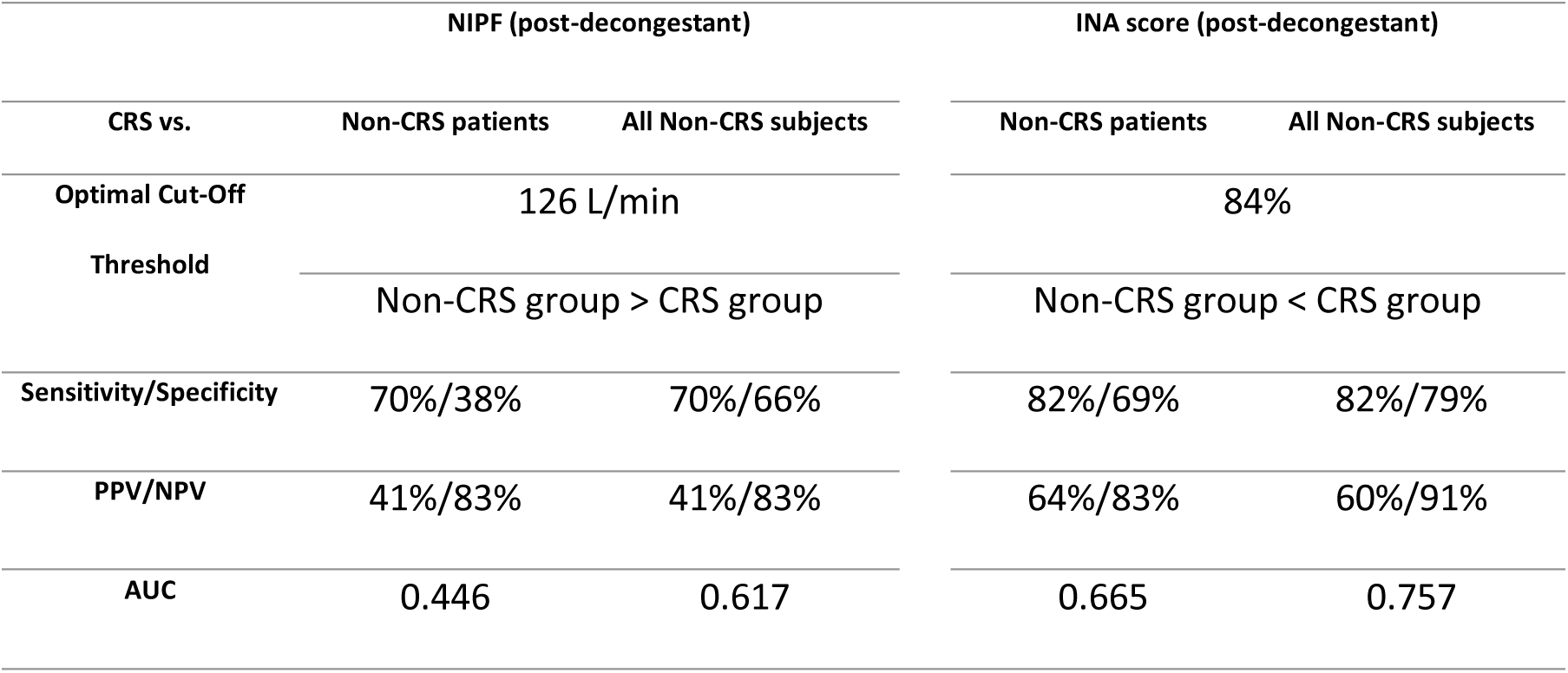
Distinguishing between subjects with CRS and subjects without based on the post-decongestant percentage change.

**Fig.5.**
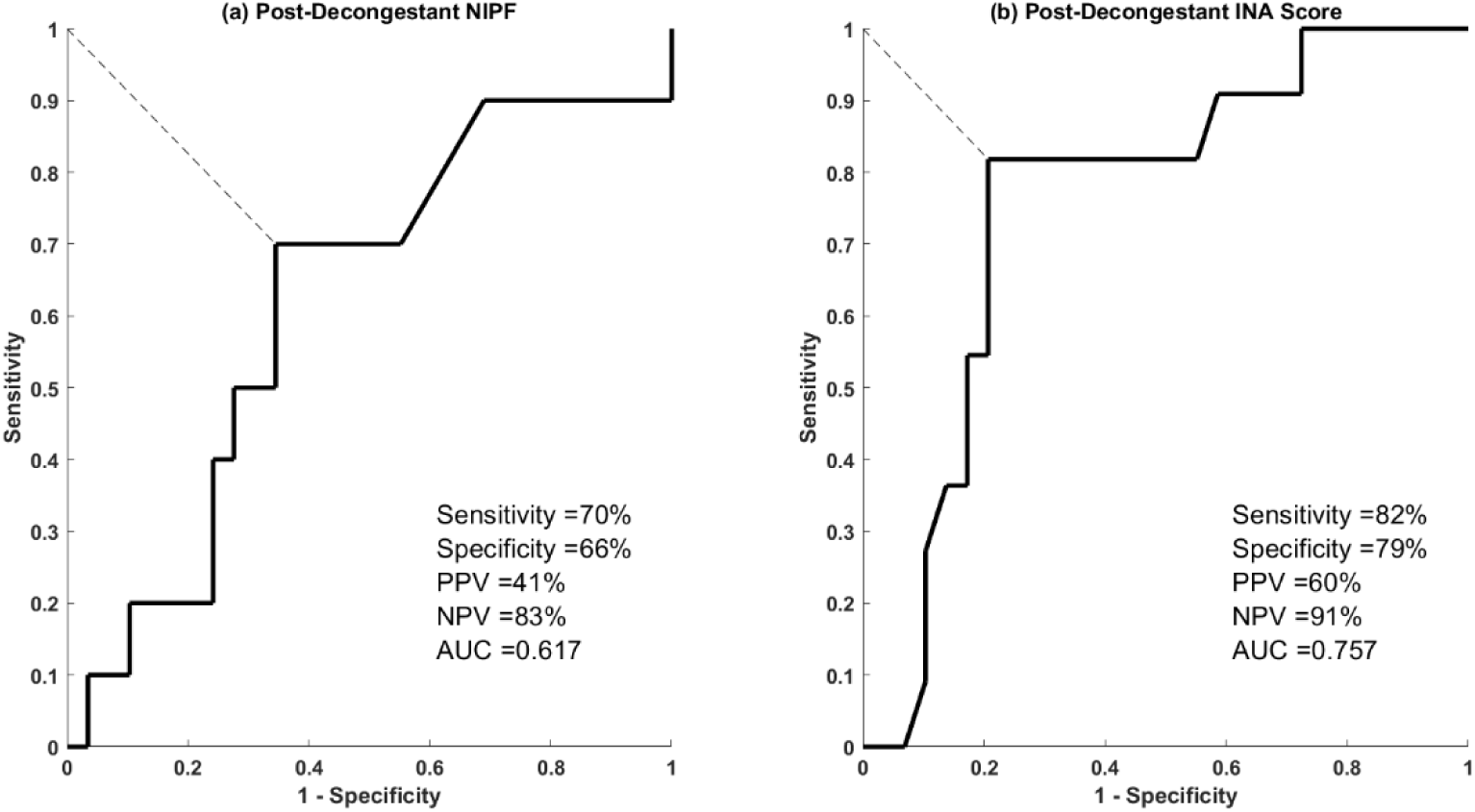
ROC curve for post-decongestant measurements in (a) NIPF and (b) INA score.

## Discussion

In terms of distinguishing between CRS patients and all non-CRS subjects (both patients and controls together), the results demonstrated that the INA score was superior to NIPF in terms of sensitivity (82% vs. 70%), specificity (79% vs. 66%), PPV (60% vs. 41%), NPV (91% vs. 83%). When controls were excluded and only the CRS and non-CRS patients were compared, the specificity for NIPF dropped sharply to 38% whilst for NAD it remained relatively high at 69%. This suggest that NIPF becomes an ineffective diagnostic tool when attempting to detect CRS from patients with other causes of nasal blockage whilst the NAD remains an effective diagnostic tool.

The need to introduce post decongestant measurements in this technique was because there was a wide variation between normal and abnormal congestion found both in controls and patients mainly due to the natural nasal cycle [28]. and nasal decongestion eliminates this confounding factor. This is especially true for INA scores whereby the pre-decongestant values were all very similar across all subject groups and the differences between groups were only apparent following nasal decongestant.

For NIPF, improvements in the nasal airflow, following decongestant, were shown simply through a higher flow measurement. Whereas in the case of the INA score, an improvement in nasal airflow is paradoxically indicated by a decreased INA score because the acoustic power is being spread over a wider range of frequencies, thus diluting the INA score. This finding additionally demonstrates that the NAD could be used for monitoring nasal airway patency, similarly to NIPF.

Whilst post-decongestant change in the measured values have been described as a way to distinguish between rhinitic and structural obstructions in the nose [29], we have shown that the post-decongestant effect could also be used to distinguish between different types of rhinitic conditions such as allergic rhinitis and CRS. The results in this study demonstrate a difference between the CRS group and the non-CRS groups, with the latter generally having higher post-decongestant percentage changes in both the NIPF and INA score (post-decongestant increase for the former, decrease for the latter). This was further demonstrated by the fact that only the controls and non-CRS patients showed statistically significant percentage changes for both NIPF and NAD whilst in CRS patients they were not statistically significant for either measurement methods. In the case of NAD this was despite the existence of a few very extreme outliers in both of the non-CRS groups which would have affected the statistical test. It must be noted that non-allergic rhinitis patients were not included in this study so as to minimise the heterogeneity of our pilot study group however this will be further explored in future studies.

Rhinitis causes a nasal obstruction through turbinate enlargement. However, there are different mechanisms which causes turbinate enlargement. Allergic rhinitis is caused by increasing congestion of the venous sinuses which can be reversed with the use of decongestants. In the case of rhinitis caused by a more chronic condition, such as in CRS, there may be more tissue inflammation as opposed to venous congestion making the enlargement less likely to be reversed through pure medications which may necessitate surgery [30]. This outcome is in parallel with CRS predominantly affecting the middle turbinate and sinuses, whereas AR largely affects the inferior turbinate. It could therefore be argued that the inferior turbinate being the larger structure and more involved with the nasal vascular regulated nasal cycle will also result in greater post-decongestant changes. Conversely, bilateral NIPF has been shown to significantly increase following functional endoscopic sinus surgery (FESS) in addition to correlating with improved quality of life in patients [31] which would imply that more drastic treatment regimens would be required to see a change in NIPF in CRS. Another study did suggest that CRS patients may require a more long term course of decongestants to be effective, compared to allergic rhinitis [32].

In the context of only applying one single on-the-day dose of decongestant to the subjects, CRS and DNS can both be considered non-decongestable conditions (the latter being totally non-decongestable). This meant that the difference in the results between CRS and non-CRS subjects may be due to DNS coincidently being more prevalent in the former group. However, as shown in Table 1, the majority of patients from both groups have DNS which suggests that the condition may not be the deciding factor.

Fundamentally, nasal breathing sounds are caused by the turbulence of air as it is breathed into and out of the nose [22]. Such sounds are affected by the internal structure and tissue characteristics within the nose. [22]. However, when a sensor is placed in the path of the nasal airflow, the expired air hitting the microphone will generate additional acoustic signal, making the overall signal during the expiration phase of breathing much greater [23]. As shown in Alshaer et al [23] and Primov-Fever *et al*’s studies [26] the expiration sounds appear so dominant that the inspiratory sounds can appear insignificant in comparison. For the NAD it was desired that only the sound of airflow-airway interactions should be detected. Hence, the NAD functions less like an airflow meter and more like an electronic stethoscope that detects specific features akin to listening to heart sounds. Furthermore, such placements of the microphones enable equal sensitivities to both inspirations and expirations.

The NAD can measure not only the breathing sounds from both sides of the nose unilaterally but also simultaneously; allowing for a more natural breathing rhythm, as well as measuring both inspirations and expirations. More importantly, the NAD can be further modified so that it also has a computer read out which enables the patient to see their nasal breathing in real time which could visually reinforce their breathing disorder. As such the NAD is envisaged as a much more versatile tool capable of a wide range of different assessments.

This pilot study has demonstrated the significant potential of the NAD in diagnosing CRS, one of the most common and burdensome of nasal conditions in addition to being able to measure improved nasal patency following treatment. Furthermore, the INA score is only one of many possible metrics which can be derived from nasal breathing sounds and other metrics could also be explored such as expiration sounds or using the NAD to diagnose other conditions such as a DNS.

## Acknowledgments

The authors would like to thank the subjects and the hospital nurses for their assistance in the study.

